# Chimeras of Kinesin-6 and Kinesin-14 reveal head-neck-tail domain functions and dysfunctions that lead to aneuploidy in fission yeast

**DOI:** 10.1101/2024.08.15.608119

**Authors:** Priyanka Sasmal, Makito Miyazaki, Frédérique Carlier-Grynkorn, Phong T. Tran

## Abstract

Kinesin motors play diverse roles in cells, including spindle assembly and chromosome segregation. Each kinesin has three general domains – the motor head, neck, and tail. As microtubule (MT) motors, kinesins have directionality, walking toward the plus- or minus-end of a MT. Plus-end kinesins have their motor head at the N-terminus, while minus-end kinesins have their motor head at the C-terminus. Interestingly, *in vitro* data indicate that the motor head does not dictate directionality. Here, we seek to understand the cellular function of each kinesin domain. We systematically created chimeras of fission yeast kinesin-6 Klp9 (a plus-end kinesin localized at the spindle midzone to slide the MTs and elongate the spindle) and kinesin-14 Pkl1 (a minus-end kinesin localized at the spindle poles to focus MTs). Our *in vivo* data reveal that the tail dictates cellular localization, and in some cases directionality of the motor head; the motor head produces binding and sliding forces affecting spindle function; and the neck modulates the forces of the motor head. Specifically, Pkl1-head, when put on Klp9-neck-tail, walks toward the spindle midzone and slides MTs faster than the wild-type Klp9. This results in spindle breakage and aneuploidy. In contrast, Klp9-head, when put on Pkl1-neck-tail, localizes to the spindle poles, but failed to properly focus MTs, leading to abnormal MT protrusions. This results in asymmetric displacement of the chromosomes and aneuploidy. Our studies reveal domain-dependent control of motor localization, direction, and force production, whose dysfunctions lead to different modes of aneuploidy.

## Introduction

Kinesins are molecular motors performing diverse roles in cells, ranging from vesicle transport to spindle assembly and chromosome segregation ^1,2^. There are currently ~45 kinesins grouped into 14 families, with cellular function uniquely attributed to each kinesin family ^3,4^. In general, kinesin domain organization consists of the motor head (h), neck (n), and tail (t), with only the heads showing high sequence homology among the different kinesins ^5,6^. Kinesins bind microtubules (MTs) via the motor head, and coupled to ATP-hydrolysis, can directionally walk toward either the MT plus- or minus-end, and/or polymerize/depolymerize MTs ^1,2^. Kinesin tail is involved in oligomerization, making the kinesin a dimer or tetramer ^7,8^. Kinesin neck modulates the activity of the motor head ^9–11^. Interestingly, plus-end directed kinesins have their heads at the N-terminus, and minus-end directed kinesins have their heads at the C-terminus ^3,4^. Surprisingly, *in vitro* motility assays of recombinant chimeric kinesins, where plus- and minus-end heads were swapped, revealed that the head does not dictate motor directionality ^9^. Instead, truncations or mutations in the neck reversed motor directionality ^12–14^. We seek to extend these findings *in vivo*, to determine precise domain functions of kinesins and to reveal the biological consequences of altered domain properties.

The fission yeast *Schizosaccharomyces pombe* has been a useful model organism to investigate kinesins. Specifically, kinesin-6/Klp9 ^15–19^ and kinesin-14/Pkl1 ^20–26^ have been extensively characterized for their distinct and evolutionarily conserved roles in mitosis. Klp9 is a plus-end directed tetrameric motor that localizes to the anaphase spindle midzone, to slide antiparallel MTs apart, thus elongating the spindle and separating the sister chromosomes ^15–19^. The absence of Klp9 results in reduced spindle elongation rate, prolonged mitosis, and increased DNA damage ^15,27^. Pkl1 is a minus-end directed motor that localizes to the spindle poles to focus pole MTs ^21–25^. The absence of Pkl1 results in abnormal pole MT protrusions and misplaced chromosomes at the end of mitosis ^23,25^. These two motors are thus different in localization, directionality, and function, and therefore are ideal to dissect the function of individual kinesin domains through chimeras. By systematically swapping the head, neck, and tail of Pkl1 into Klp9, and vice versa, to create kinesin chimeras, we show that the tail domain, and not the neck, controls motor localization and directionality in cells. The neck modulates the motor head activity. Further, we reveal chimeric kinesin domain dysfunctions that lead to spindle assembly defects, chromosome segregation error, and aneuploidy.

## Results

In fission yeast, the gene encoding plus-end directed kinesin-6/Klp9 (K9) is located on chromosome II, with its motor head at the N-terminus; and the gene encoding minus-end directed kinesin-14/Pkl1 (P1) is located on chromosome I, with its motor head at the C-terminus. We delineated the head (h), neck (n) and tail (t) domain of these kinesins using a combination of SMART, COILED-COILS, and AlphaFold2 predictive applications available on PomBase (www.pombase.org). The head contains the ATPase globular domain; the tail begins at the first coiled-coil region; and the short neck links the head to the tail (Fig 1A). We then created a series of K9 chimeras by systematically replacing K9 head, neck, and tail with P1 head, neck, and tail, respectively. Each chimera was expressed at the K9 locus under the K9 promotor (Fig. 1A). For example, Klp9 wild-type is K9hnt (Klp9 head-neck-tail), Klp9 chimera with Pkl1 head is P1h:K9nt (Pkl1 head:Klp9 neck-tail), etc. We also created a complementary series of P1 chimeras, with K9 head, neck, and tail respective replacement, and expressed at the P1 locus under the P1 promotor (Fig. 1A). All motors were tagged with eGFP for visualization. Cells also expressed mCherry-Atb2 (tubulin) for MT visualization.

**Figure 1.**
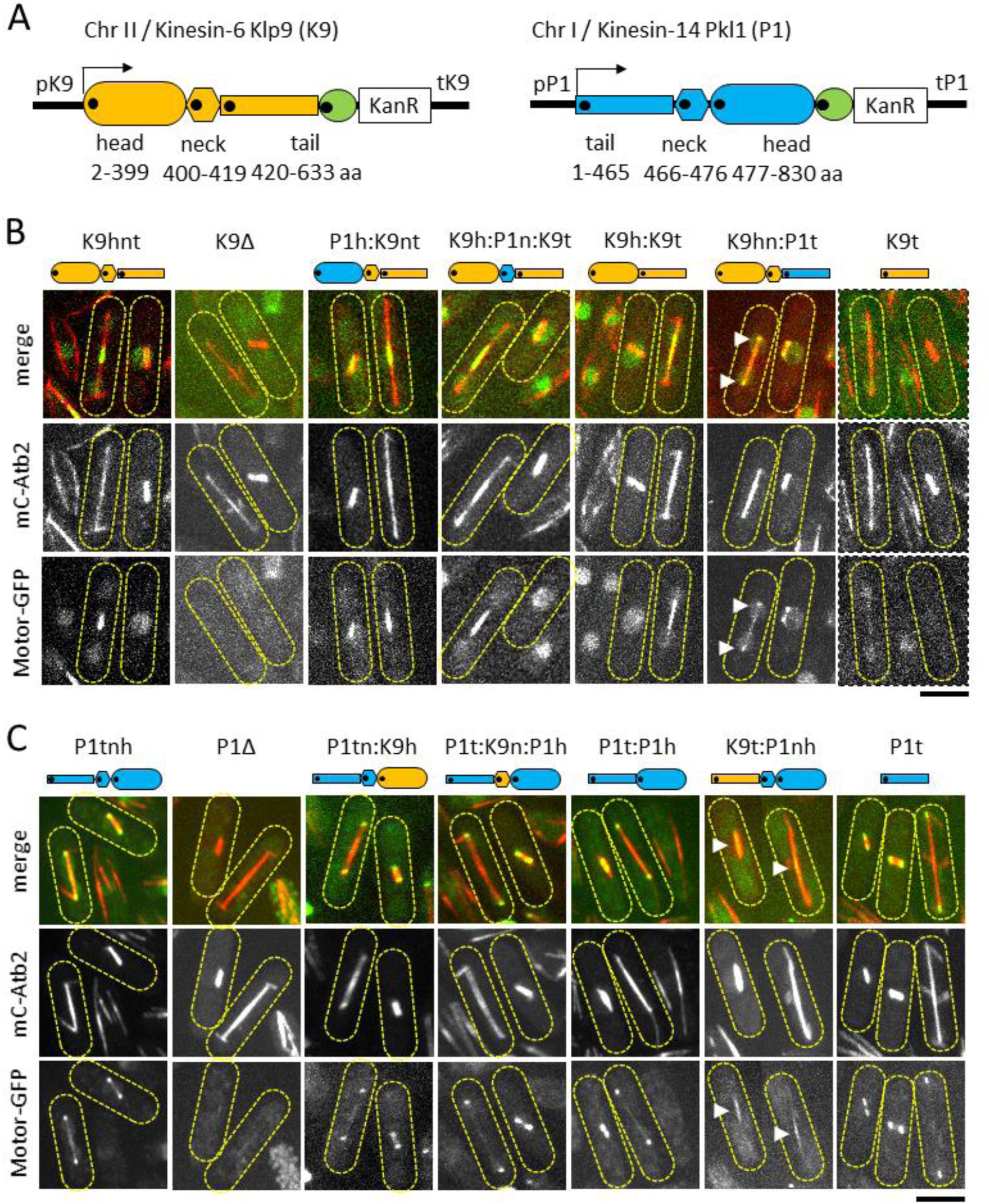
Kinesin tail domain determines motor localization on spindles. **A.** Cartoons showing the head, neck, and tail domains of fission yeast wild-type kinesin-6 Klp9 (orange) and kinesin-14 Pkl1 (blue), detailing their amino-acid length and chromosomal location. Green circles represent eGFP tagged at the C-terminus of each motor. Black dot indicates the N-terminus of each domain. pK9/tK9 and pP1/tP1 indicate the promotor and terminator of the respective motor. KanR is the selection marker. **B.** Images of the Klp9 series. Merge color and individual black-white fluorescence of mCherry-Atb2 (tubulin) and motor-eGFP (motors), of mitotic cells (yellow dashed outline) with short metaphase spindles and long anaphase spindles. The chimeric and truncated motors are labelled as cartoons and text. White arrow heads highlight the localization of the chimeric motor K9hn:P1t on the spindle. Scale bar, 5 µm. **C.** Images of the Pkl1 series. Merge color and individual black-white fluorescence of mCherry-Atb2 (tubulin) and motor-eGFP (motors), of mitotic cells (yellow dashed outline) with short metaphase spindles and long anaphase spindles. The chimeric and truncated motors are labelled as cartoons and text. White arrow heads highlight the localization of the chimeric motor K9t:P1nh on the spindle. Scale bar, 5 µm.

### The tail dictates localization of chimeric kinesin motors on the spindle MTs

As a plus-end directed kinesin, the wild-type Klp9 (K9hnt) localized to the anaphase spindle midzone (Fig. 1B), which contains antiparallel plus end MTs, consistent with previous reports for this motor ^15–17,19^. As a minus-end directed kinesin, the wild-type Pkl1 (P1tnh) localized to the spindle poles (Fig. 1C), which contain minus end MTs, also consistent with previous findings for this motor ^21,23,25^.

For the Klp9 chimeric series, chimera P1h:K9nt (which has the P1 minus-end directed head), chimera K9h:P1n:K9t (which has the P1 neck), and truncation K9h:k9t (which has no neck), all localized to the spindle midzone (Fig. 1B), similar to wild-type Klp9. In contrast, chimera K9hn:P1t (which has the P1 tail) localized to the spindle poles (Fig. 1B), similar to wild-type Pkl1 (Fig. 1C). Truncations K9h and K9hn (which has no tails), showed no spindle localization (Fig. 1SA). In parallel, for the Pkl1 chimeric series, chimera P1tn:K9h (which has the K9 plus-end directed head), chimera P1t:K9n:P1h (which has the K9 neck), and truncation P1t:P1h (which has no neck), all localized to the spindle poles (Fig. 1C), similar to the wild-type Pkl1. In contrast, chimera K9t:P1nh (which has the K9 tail), localized to the spindle midzone (Fig. 1C), similar to the wild-type Klp9 (Fig. 1B). Truncations P1h and P1nh showed no spindle localization (Fig. 1SB).

The results thus far highlight kinesin tail as the determinant of motor localization; and suggests that the motor head and neck do not determine motor localization. Interestingly, the truncated K9t did not localize to the spindle midzone (Fig. 1B), but the truncated P1t localized to the spindle poles (Fig. 1C). This result further suggests that Klp9 tail may control motor head position through controlling the head directionality; and Pkl1 tail may control motor position via directly binding to the spindle poles.

### Kinesin neck positively modulates the motor head activity

We next analyzed spindle dynamics of the Klp9 chimeric series through time-lapsed live-cell imaging. Fission yeast undergoes 3-phase spindle elongation (1 = prophase; 2 = metaphase-anaphase A; 3 = anaphase B) ^28^, with a dramatic Klp9-dependent length increase during phase-3/anaphase B ^15,16,19^. Imaging revealed that all chimeras performed 3-phase spindle elongation (Fig. 2A), with similar phase-1 and phase-2 velocities (Fig. 2B), but distinct and varied phase-3 velocities (Fig. 2B). Klp9-deletion (K9Δ), Klp9-truncations (K9h:K9t, K9t), chimeras K9h:P1n:K9t and K9hn:P1t all had significantly lower anaphase spindle sliding velocities compared to the wild-type Klp9 (K9hnt) (Fig. 2B, 2C). Interestingly, while K9Δ exhibited ~50% slower velocities compared to wild-type K9hnt (Klp9 0.68 µm/min versus K9Δ 0.35 µm/min), consistent with previous findings ^15,16,19^, the no neck truncation K9h:K9t and the swapped neck chimera K9h:P1n:K9t had only slightly slower anaphase velocities (~10-15% slower) than the wild-type K9hnt (K9h:P1n:K9t 0.62 µm/min and K9h:K9t 0.58 µm/min) (Fig. 2B, 2C). This result indicates that the absence of the neck does not abolish motor head activity; instead, the neck domain positively modulates the motor head activity.

**Figure 2.**
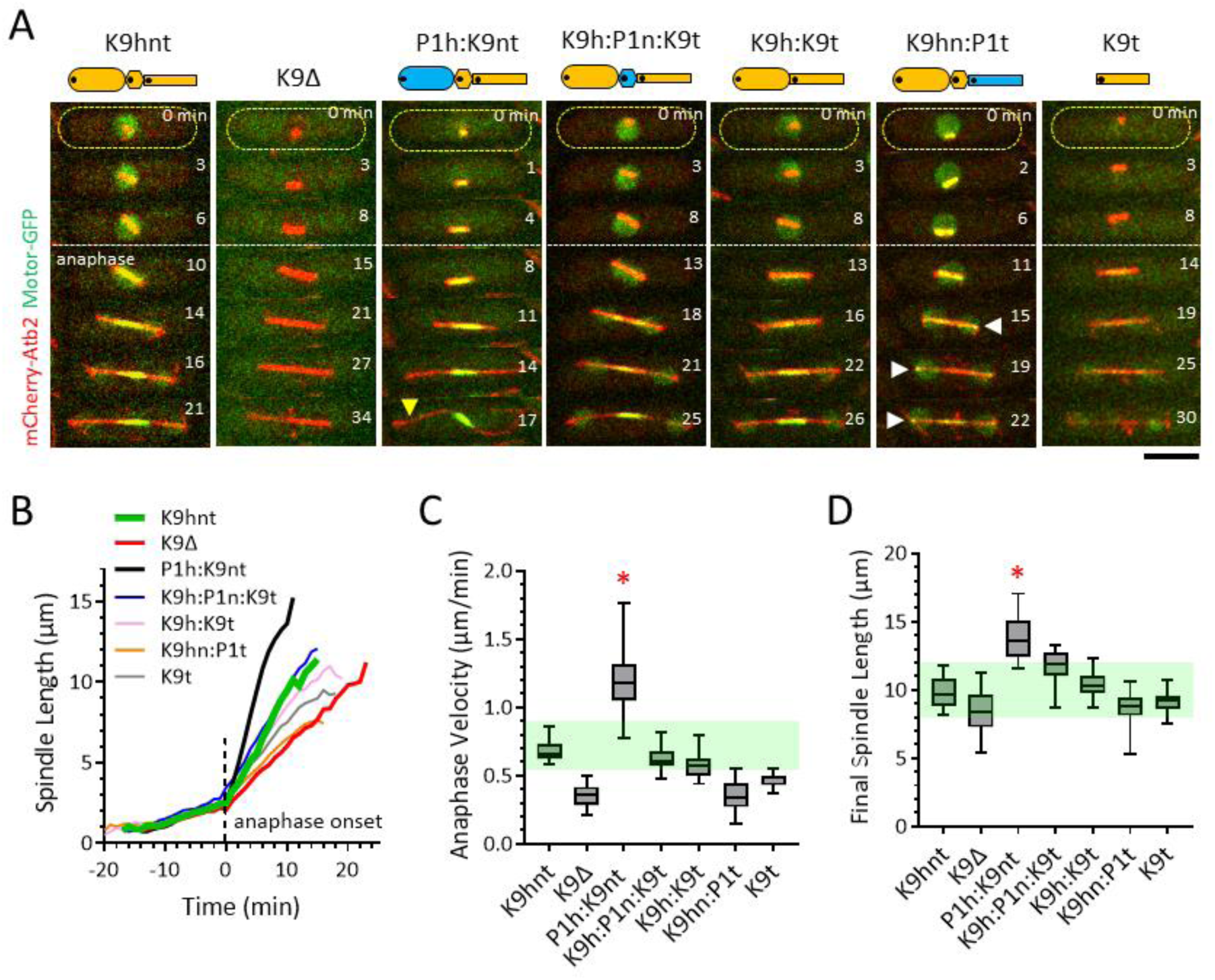
Chimera P1h:K9nt elongates the spindle faster than wild-type Klp9. **A.** Time-lapsed images of spindle dynamics throughout mitosis for the Klp9 series. Cells expressed mCherry-Atb2 (tubulin) and motor-eGFP (motors). Scale bar, 5 µm. **B.** Plot of Mean Spindle Length versus time throughout mitosis for the Klp9 series. For each strain, 20 cells were measured to derive the mean. **C.** Box plot of Anaphase Velocity for the Klp9 series. Mean ± Standard Deviation, t-test p-values with respect to wild-type: K9hnt 0.68 ± 0.07 µm/min (n=20); K9Δ 0.35 ± 0.08 µm/min (n=20, p<0.001); P1h:K9nt 1.19 ± 0.24 µm/min (n=20, p<0.001); K9h:P1n:K9t 0.62 ± 0.08 µm/min (n=20, p<0.03); K9h:K9t 0.58 ± 0.10 µm/min (n=20, p<0.001); K9hn:P1t 0.35 ± 0.11 µm/min (n=20, p<0.001); K9t 0.45 ± 0.05 µm/min (n=20, p<0.001). **D.** Box plot of Final Spindle Length at end of anaphase for the Klp9 series. Mean ± Standard Deviation, t-test p-values with respect to wild-type: K9hnt 9.75 ± 1.11 µm (n=20); K9Δ 8.38 ± 1.59 µm (n=20, p<0.005); P1h:K9nt 13.80 ± 1.61 µm (n=20, p<0.001); K9h:P1n:K9t 11.74 ± 1.17 µm (n=20, p<0.001); K9h:K9t 10.37 ± 1.02 µm (n=20, p=0.07); K9hn:P1t 8.66 ± 1.12 µm (n=20, p<0.005); K9t 9.16 ± 0.73 µm (n=20, p=0.06).

### Chimera P1h:K9nt elongates the spindle faster than wild-type Klp9

Surprisingly, the chimera P1h:K9nt showed faster spindle velocity compared to the wild-type Klp9 (Fig. 2A, 2B). Whereas wild-type K9hnt attained spindle velocity of 0.68 µm/min, the chimera P1h:K9nt reached spindle velocity of 1.19 µm/min, or ~1.5x faster (Fig. 2C). As strains K9hnt and P1h:K9nt have similar cell lengths at mitosis (Fig. 2SA), similar mitosis duration (Fig. 2SB), and similar anaphase duration (Fig. 2SC), the faster spindle velocity of P1h:K9nt therefore resulted in an abnormally long final spindle length (Fig. 2D), that manifested as S-shaped spindles due to spindle buckling upon reaching the cell tip cortex (Fig. 2A, yellow arrow head, t = 17 min).

We next probed a mechanism for the observed P1h:K9nt chimera-dependent fast spindle elongation. Fission yeast anaphase spindle elongation velocity scales with the number of Klp9 motor at the midzone – more Klp9, faster speed, less Klp9, slower speed ^16^. We thus measured signal intensities, as a proxy for the number of motors, on 10-µm long anaphase spindles (Fig. 3SA). Line-scan analysis revealed that the wild-type K9hnt and the chimera P1h:K9nt have nearly identical signal intensity profiles (Fig. 3A), with the same total signal (Fig. 3SB) and the same signal at the midzone (Fig. 3SC). This result indicates that the ~1.5x faster than wild-type spindle elongation speed of chimera P1h:K9nt is not a consequence of an increase in the number of P1h:K9nt, but is likely due to the inherent faster MT sliding speed of the Pkl1 motor head.

**Figure 3.**
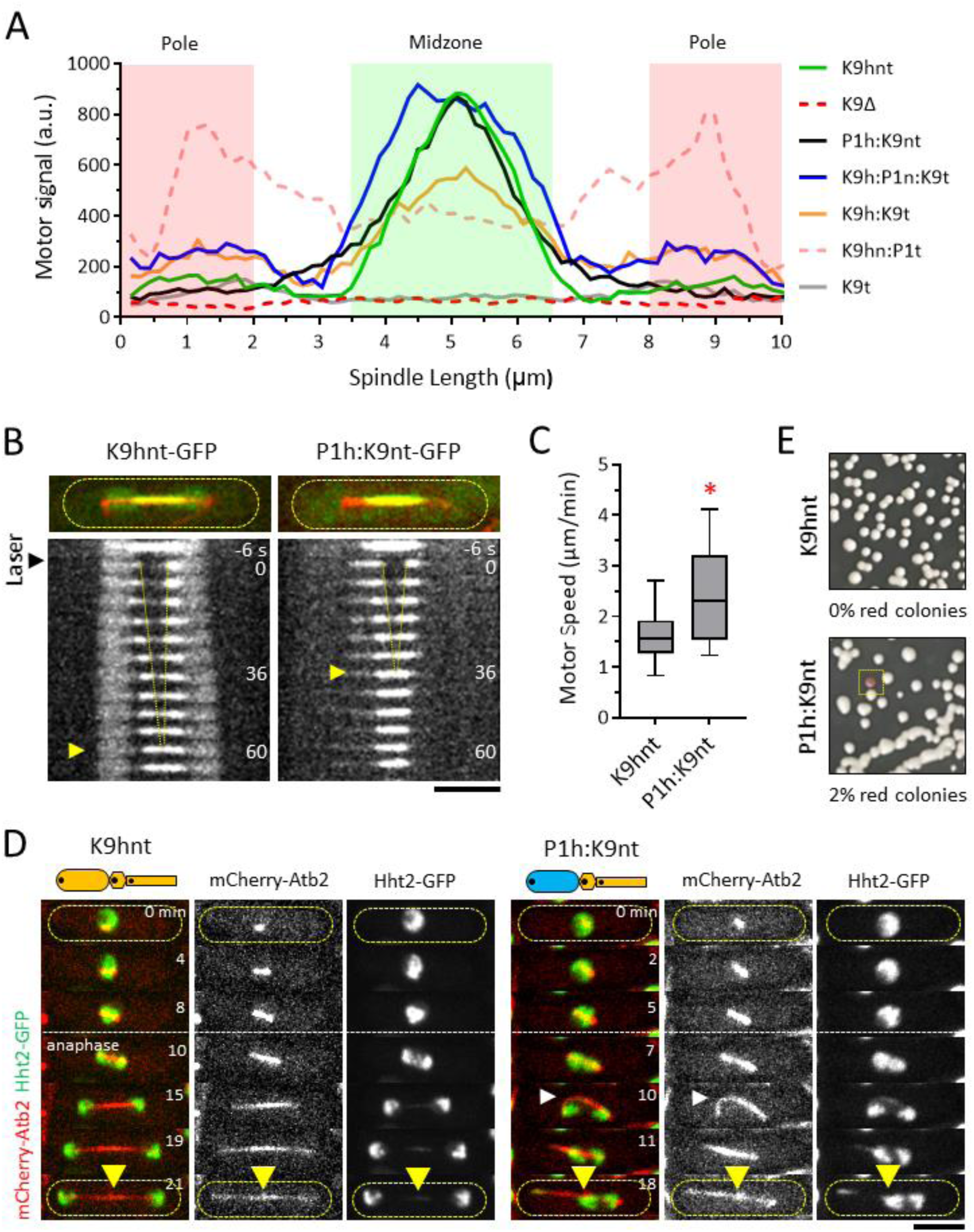
The inherent faster speed of P1h:K9nt results in spindle breakage and aneuploidy. **A.** Line scan plot of mean Motor Signal intensity on 10-µm long spindles for the Klp9 series. For each strain, 20 cells were measured to derive the mean. **B.** Time-lapsed images of photo-bleaching recovery of wild-type Klp9-GFP and chimera P1h:K9nt-GFP. Yellow arrow marks time point when fluorescent signal completely recovered after being photo-bleached. Scale bar, 5 µm. **C.** Box plot of Motor Speed, a proxy of time to complete recovery, for wild-type Klp9 and chimera P1h:K9nt. Mean ± Standard Deviation, t-test p-values: K9hnt 1.62 ± 0.48 µm/min (n=33); P1h:K9nt 2.39 ± 0.92 min (n=29, p<0.001). **D.** Time-lapsed images of spindle dynamics throughout mitosis for wild-type Klp9 and chimera P1h:K9nt. Cells expressed mCherry-Atb2 (tubulin) and Hht2-GFP (histone). White arrow head indicates a P1h:K9nt buckled spindle that subsequently breaks (time = 10 min and 11 min). Yellow arrow head indicates medial position where cell divides. Scale bar, 5 µm. **E.** Images of artificial minichromosome loss assay for wild-type Klp9 and chimera P1h:K9nt. White colonies indicate normal cell division and no artificial minichromosome loss. Red colonies (square box) indicate loss of the artificial minichromosome. Wild-type Klp9 has zero red colonies (n=753 colonies). P1h:K9nt has 2% red colonies (27 red, n=1447 colonies).

Indeed, in a different set of experiments where we photo-bleached the center of the spindle midzone, we observed ~1.5x faster directed recovery of chimera P1h:K9nt into the bleached region compared to wild-type K9hnt (Fig. 3B). Wild-type Klp9 took 47 ± 18 sec (n=33) compared to P1h:K9nt 29 ± 6 sec (n=29) to fill-up the similar-sized bleached midzone region (Fig. 3B). The recovery signal moved sequentially in a plus-end directed manner toward the spindle center (Fig. 3B), with wild-type K9hnt speed at 1.62 µm/min and P1h:K9nt speed at 2.39 µm/min (Fig. 3C).

Together, the results indicate that the Pkl1 motor head, which was part of the minus-end directed kinesin, is moving in a plus-end directed manner when fused to the neck and tail domains of the plus-end directed Klp9. The chimera P1h:K9nt localized to the spindle midzone to slide antiparallel MTs apart and elongate the spindle, even faster than the wild-type Klp9. Thus, Klp9 tail appears to control the directionality of the motor head.

### Abnormally fast spindle elongation results in aneuploidy via spindle breakage

We examined the consequences of the abnormally fast spindle elongation velocities exhibited by the chimera P1h:K9nt. Live-cell imaging of cells expressing the chromosome marker Hht2-GFP (histone) and mCherry-Atb2 (tubulin) revealed that for the wild-type K9hnt, sister chromosomes were well-separated toward opposite cell tips by the relatively straight anaphase spindle prior to cytokinesis (Fig. 3D), resulting in two daughter cells with identical chromosomal mass or nuclei. Surprisingly, the S-shaped spindles exhibited by the majority of P1h:K9nt chimera also showed well-separated sister chromosomes (Fig. 3SE), without chromosome segregation defects at cytokinesis. In contrast, in ~2% of mitotic P1h:K9nt chimera (5 of 188 mitotic cells), the abnormally fast spindle elongation speed resulted in spindle buckling and breakage at anaphase B (Fig. 3D, white arrow head, t = 10 min), leading to the sister chromosome masses not being well-separated at cytokinesis (Fig. 3D, yellow arrow head), thus creating aneuploidy subsequent to cell division. This novel aneuploidy phenotype suggests that the rate of spindle elongation at anaphase, defined by the motor head sliding activity, is regulated to prevent chromosome segregation error. Dysfunction in the motor head, such as faster than normal sliding activity, can cause spindle breakage and aneuploidy.

We confirmed the aneuploidy phenotype of the P1h:K9nt chimera through the robust artificial mini-chromosome loss assay ^29^. An artificial mini-chromosome was introduced into the wild-type K9hnt and chimera P1h:K9nt strain. Loss of the mini-chromosome turns the fission yeast from white to red, signifying chromosome segregation errors. We observed zero red colonies for the wild-type K9hnt, and ~2% red colonies in the chimera P1h:K9nt (Fig. 3E). This result is identical to the ~2% frequency of spindle breakage observed in chimera P1h:K9nt (Fig. 3D), thus confirming that spindle buckling and breakage due to abnormally high rate of spindle elongation at anaphase can cause chromosome segregation error and aneuploidy.

### Pkl1 chimeras do not prevent abnormal MT protrusions

Turning to the Pkl1 series, we examined abnormal MT protrusions at the spindle poles. Pkl1 localizes to the spindle poles to focus MTs; and the absence of Pkl1 (P1Δ) results in abnormally long pole MT protrusions ^21,23,25^. We observed zero MT protrusions in wild-type Pkl1 cells (Fig. 4A, 4B); and ~12% of spindles with abnormal pole MT protrusions of 5-µm or longer in Pkl1-deletion (P1Δ) (Fig. 4A, yellow arrow head), consistent with previous results ^23^. Surprisingly, all chimeras and truncated strains in the P1 series exhibited abnormally long MT protrusions at frequencies similar to P1Δ (Fig. 4A, 4B). The result indicates the chimeras P1tn:K9h (which has Klp9 head), P1h:K9n:P1h (which has Klp9 neck), and truncation P1t:P1h (which has no neck), while localized correctly to the spindle poles, cannot focus MTs to prevent abnormal protrusions.

**Figure 4.**
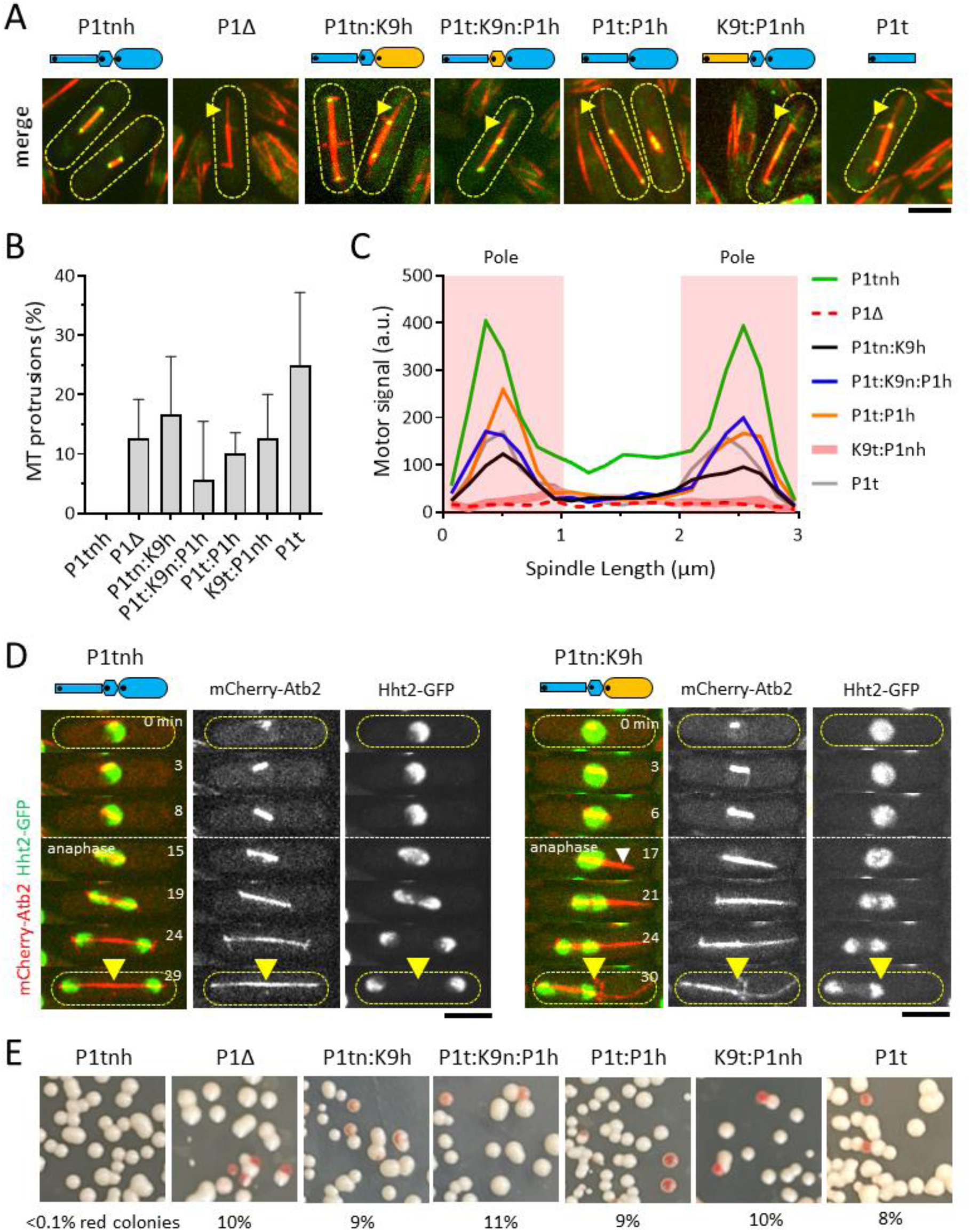
Abnormally long spindle pole MT protrusions results in nuclear mis-position and aneuploidy. **A.** Images of the abnormally long pole MT protrusions present in the Pkl1 series. Yellow arrow head highlights the MT protrusion. Scale bar, 5 µm. **B.** Bar plot of frequency of mitotic cells with abnormally long MT protrusions ≥ 5-µm. Mean ± Standard Deviation, t-test p-values with respect to P1Δ: P1tnh 0% (zero protrusions, n>100 spindles); P1Δ 12 ± 7% (10 protrusions, n=76 spindles); P1tn:K9h 16 ± 10% (9 protrusions, n=56 spindles, p=0.41); P1t:K9n:P1h 6 ± 10% (2 protrusions, n=34 spindles, p=0.14); P1t:P1h 10 ± 4% (5 protrusions, n=52 spindles, p=0.65); K9t:P1nh 13 ± 7% (9 protrusions, n=75 spindles, p=0.83); K9t 25 ± 12% (8 protrusions, n=31 spindles, p=0.21). **C.** Line scan Plot of mean Motor Signal intensity on 3-µm long spindles for the Pkl1 series. **D.** Time-lapsed images of spindle dynamics throughout mitosis for wild-type Pkl1 and chimera P1tn:K9h. Cells expressed mCherry-Atb2 (tubulin) and Hht2-GFP (histone). White arrow head indicates a P1tn:K9h MT protrusion (time = 17 min) that pushed the chromosome mass to the cell center, where subsequent cell division (yellow arrow head) resulted in aneuploidy. Scale bar, 5 µm. **E.** Images of artificial minichromosome loss assay for the Pkl1 series. White colonies indicate normal cell division and no artificial minichromosome loss. Red colonies indicate loss of the artificial minichromosome. Wild-type Pkl1 (1 red, n=1301 colonies); P1Δ (149 red, n=1639 colonies); P1tn:K9h (131 red, n=1511 colonies); P1t:K9n:P1h (172 red, n=1459 colonies); P1t:P1h (135 red, n=1582 colonies); K9t:P1nh (208 red, n=2013 colonies); K9t (177 red, n=2206 colonies).

To probe a potential reason for why chimeras cannot prevent abnormal MT protrusions, we measured motor signal intensities, as a proxy for the number of motors, on 3-µm long spindles (Fig. 4SA). Line-scan analysis revealed that all Pkl1 chimeras and truncations have significantly less signal at the spindle poles compared to the wild-type P1tnh (Fig. 4C, Fig. 4SB, 4SC). Indeed, the Pkl1-tail (P1t) alone can localize to the spindle poles (Fig. 4A), albeit to a lesser extent than the wild-type P1tnh (Fig. 4C). All other chimeras also localized to the spindle poles, with similar signal to P1t (Fig. 4C). The result provides an explanation for why the chimera P1tn:K9h, and all other Pkl1 chimeric and truncated strains, cannot prevents abnormally long MT protrusions: there may not be enough motors at the poles to prevent protrusions. Consistent with this hypothesis, it was previously shown that the amount of Pkl1 motor localized at the spindle pole is critical for MT minus-end focusing ^25,26^.

### Abnormally long spindle MT protrusions result in aneuploidy via misplaced daughter nuclei

We previously showed that Pkl1-deletion dependent abnormally long spindle MT protrusions caused aneuploidy due to misplaced daughter nuclei at cell division ^23^, i.e., the long MT protrusions push the separated nuclei from the cell tip back to the cell center, resulting in chromosome “cut” at cytokinesis. We thus examined the consequences of the abnormally long MT protrusions in the Pkl1 chimeras. Live-cell imaging of cells expressing the chromosome marker Hht2-GFP (histone) and mCherry-Atb2 (tubulin) revealed that for the wild-type P1tnh, no pole MT protrusions was observed (Fig. 4A, 4B), and sister chromosomes were well-separated toward opposite cell tips by the relatively straight anaphase spindle prior to cytokinesis (Fig. 4D), resulting in two daughter cells with identical chromosomal mass. In contrast, we observed in Pkl1-deletion (P1Δ) ~12% long MT protrusions greater than 5-µm emanating from a spindle pole at anaphase(Fig. 4A, 4B), to push the divided chromosome mass to the cell center (Fig. 4SD, white arrow head, t = 12 min), where subsequent cytokinesis will create aneuploid cells (Fig. 4SD, white arrow head, t = 25 min). Similarly, in the various Pkl1 chimeras, we observed ~10-16% of mitotic cells with the same phenotype of abnormally long spindle MT protrusions (Fig. 4D, white arrow head, t = 17 min), and resulting chromosome misplacement (Fig. 4D, yellow arrow head, t = 30 min), and subsequent aneuploidy.

We confirmed the aneuploidy phenotype of the Pkl1 series chimeras and truncations through the artificial mini-chromosome loss assay ^29^. We observed less than 0.1% red colonies for the wild-type P1tnh, and between ~8-11% red colonies in the Pkl1 series of chimeric and truncated motors (Fig. 4F). This is similar to the frequency of aneuploidy observed due to abnormally long MT protrusions at spindle poles (Fig. 4B). The result confirmed that abnormally long MT protrusions from the spindle poles due to dysfunction of Pkl1 localization at the spindle poles can misplace the chromosomes leading to aneuploidy at cell division.

## Discussion

Our current work with chimeric mitotic kinesin-6/Klp9 and kinesin-14/Pkl1 in *S. pombe* cells reveals functional properties of kinesin domains, and how dysfunctions in motor properties can result in errors in spindle assembly dynamics, leading chromosome segregation errors and aneuploidy. First, the motor head does not dictate positioning and directionality of the motor (Fig. 1), consistent with *in vitro* analysis ^30^. Swapping the minus-end directed Pkl1-head onto plus-end directed Klp9 does not impact Klp9 localization and directionality (Fig. 1). Second, the neck modulates the function of the head (Fig. 2), but plays no role in positioning nor directionality of the motor, contrary to *in vitro* analysis ^12,13^. Removing Klp9-neck attenuates Klp9 sliding velocity (Fig. 2), but does not impact Klp9 localization to the spindle midzone. Third, the tail appears to dictate positioning and directionality of the motor (Fig. 3). Pkl1 tail alone can localize to the spindle pole, bringing along Klp9 head; and Klp9 tail can reverse the direction of Pkl1-head, enabling it to move toward the spindle midzone. of Finally, each kinesin may be evolutionarily tuned for specific and optimal function, and altering the properties of the kinesins can result in aneuploidy, e.g., making Klp9 slide MTs faster will result in spindle breakage and aneuploidy (Fig. 3), and making Pkl1 localize less to the spindle poles will result in abnormally long MT protrusions and aneuploidy (Fig. 4).

As it stands, our current work in cells confirms some concept set forth the by *in vitro* studies ^12,13,30^, that the motor head does not itself dictate directionality; and our work reveals that in fact the tail controls motor directionality and positioning. We show that in cells the neck only modulates the function of the head, but plays no role in motor directionality, which contradicts findings from *in vitro* studies ^12,13^. One potential explanation for the discrepancy is that the *in vitro* studies were often performed with truncated motors with short or no tails. Further, there may be an alternative explanation, not involving the neck tail, for what controls motor directionality: the physical location of the motor head at either the N-terminus or the C-terminus of the amino-acid sequence defines its directionality ^13^.

One surprise is that the chimera P1h:K9nt, which put the Pkl1 head onto the Klp9 neck-tail, can slides the spindle ~1.5x faster than the wild-type K9hnt. This indicates that the Pkl1 head is inherently faster than the Klp9 head. We currently do not know the reason for the faster speed of the Pkl1 head, but surmise that its faster speed may be due to its ability to hydrolyze ATP faster ^31^, or its ability to bind MTs stronger ^32^. These parameters may be determined through *in vitro* analysis.

A second surprise is that the chimera P1tn:K9h, which put the Klp9 head onto the Pkl1 neck and tail, cannot prevent abnormal MT protrusions, even though it localizes properly to the spindle poles, albeit at less concentration. Further, all chimeric and truncated strains in the Pkl1 series similarly cannot prevent abnormal MT protrusions. We speculate that, as each fission yeast spindle pole contains ~25 MTs ^33–35^, a critical number of Pkl1 motors, close the wild-type Pkl1 signal at poles, is needed to bind and focus the MTs and prevent abnormal protrusions. Alternatively, the Klp9 head may inherently have weaker binding to MTs compared to Pkl1 head. These ideas may be tested both *in vivo* and *in vitro*.

We further note, due to the high reproducibility of Klp9-dependent anaphase spindle velocity and of Pkl1-dependent prevention of abnormal spindle MT protrusion in *S. pombe*, we can speculate that future chimeric experiments swapping domains of different species into fission yeast Klp9 and Pkl1 will provide new information on the biophysical properties of kinesin motors for detailed evolutionary comparison.

Finally, mitotic kinesins in general, and kinesin-6 and kinesin-14 in particular, have been implicated in cancer through either up or down regulation of motor numbers and motor activities ^36^. Our work reveals novel modes of aneuploidy with errors in spindle assembly dynamics that lead to chromosome segregation errors due to specific altered properties of kinesin-6/Klp9 or kinesin-14/Pkl1 in fission yeast. In the case of Klp9, P1h:K9nt (Pkl1 head on Klp9 neck-tail) dependent faster anaphase spindle elongation can result in buckled spindles that break, thus failing to separate the sister chromosome masses prior to cytokinesis. This is in contrast to the case of Pkl1, where P1tn:K9h (Klp9 head on Pkl1 neck-tail) dependent failure to focus MTs at the spindle poles can result in abnormally long MT protrusions that push the separated chromosome masses to the cell center, thus failing to separate the sister chromosome masses prior to cytokinesis.

Spindle elongation velocity in fission yeast appears highly regulated to maintain an “optimal” speed, as faster speed results in aneuploidy (this study), and slower speed results in DNA damage ^27^. Unlike human cells, fission yeast undergoes “closed” mitosis, during which the nuclear envelope does not breakdown at mitosis. In this context, chromosome segregation errors observed in cells with buckled broken spindles may be due to nuclear membrane-dependent tensile resistive forces opposing the fast elongating spindle ^37–40^, i.e., the abnormally fast spindle elongation speed uncouples spindle elongation from new nuclear membrane addition. In human cells, with an “open” mitosis and no nuclear membrane to resist spindle elongation, we speculate that abnormally fast spindle elongation could result in aneuploidy by uncoupling MT-to-chromosome attachment and the spindle assembly checkpoint. This can be tested by augmenting the speed of spindle elongation in human cells, perhaps through a kinesin-6 chimera.

Kinesin-14 spindle pole MT-focusing function appears to be well-conserved from yeast to human ^41^. Indeed, in cancer cells with supernumerary centrosomes, an up-regulation of kinesin-14 HSET keeps the multiple centrosomes focused to form bipolar spindles ^42^. Downregulation of HSET results in multipolar spindles and subsequent chromosome segregation errors and cancer cell death. Similarly, downregulation of fission yeast Pkl1, either through chimeras or truncations, results in unfocused spindle poles with abnormally long MT protrusions and subsequent chromosome segregation errors and cell death.

## Materials and Methods

### Construction of *S. pombe* chimeric and truncated kinesin strains

All used strains are isogenic to wild-type 972 and were obtained from genetic crosses, selected by random spore germination and replica on plates with appropriate drugs or supplements. All strains are listed in the Supplementary Data (Strain List). Gene deletion and tagging was performed as described previously ^43^. Briefly, a combination of SMART, COILED-COILS, and AlphaFold2 predictive applications available on PomBase (www.pombase.org) was used to define the head, neck, and tail of Klp9 and Pkl1. The full-length DNA sequences, including the 5’ promotor and 3’ terminator, of each motor were sub-cloned into parent vectors containing eGFP and the selection gene encoding Kanamycin-resistant at the C-terminus. Each domain of one motor was amplified from the parent vector via PCR, then swapped with the corresponding domain of the other motor, using the Gibson Assembly method ^44^. Each chimeric motor sequence was transformed into fission yeast at its parent genomic locus using the Li-Acetate method ^45^. All strains were checked for correctness via DNA sequencing (eurofinsgenomics). Florescent markers were introduced by mating the chimeric strains with other strains carrying the appropriate markers, then selecting for appropriate spores through drug-resistance selection.

### Live-cell microscopy

Cells were cultured in YE5S liquid overnight at 25ᵒC to reach OD_600nm_ ~0.5 prior to imaging. Live-cell imaging was conducted as previously described ^46^. Briefly, cells were mounted onto a slide containing 2% agarose with YE5S. Imaging was performed using an inverted Eclipse Ti-E microscope (Nikon) equipped with a spinning disk CSU-22 unit (Yokogawa). The microscope set-up included a Plan Apochromat 100X/1.4NA objective lens (Nikon), a PIFOC (perfect image focus) objective stepper, a Mad City Lab piezo stage, an Evolve EMCCD camera (Photometrics), and laser unit with 488 nm (100 mW) and 561 nm (100 mW) (Gataca Systems). The microscope was enclosed within a thermal box to keep stable temperatures of 25ᵒC. The microscope was controlled by MetaMorph 7 software (Molecular Devices). For imaging, the laser setting was GFP/488nm 10 mW and mCherry/561nm 5 mW, and camera setting was Bin 1 and EM-Gain 300. Z-stacks of seven planes spaced by 1-μm were acquired for each channel, with an exposure time of 100 ms each channel. Movies were captured at 1-min interval, for 60-90 min total.

### Photo-bleaching assay

Photo-bleaching was performed on the same microscope set-up as described above. Additionally, the MetaMorph software included the plugin iLAS2 for laser positioning and bleach-region control. For bleaching GFP, maximum laser power (488 nm, 100 mW) at 50 ms exposure was used. The bleach region was the focused laser beam, producing a circle of ~0.5 µm diameter onto the cell. Movies were captured at 100 ms exposure, 5-sec interval, for 5 min total with the laser setting at GFP (488nm, 10mW) and mCherry (561nm, 5mW).

### Image analysis

Image analysis was performed using MetaMorph 7. Maximum projections of Z-stacks from the movies were used for measurements. Velocity measurements were obtained by measuring the spindle length (mCherry-Atb2 signal) over successive time frames. Motor signal intensity was performed by line scan (6 pixel wide) of the average motor intensity (motor-eGFP) on the spindles. All values, including mean ± standard deviation and t-test were performed in Excel (Microsoft). Plots were generated using Prism 10 (GraphPad).

### Minichromosome loss assay

The minichromosome loss assay was performed as previously described ^29^. Briefly, the chimeric strains were mated with a strain carrying the artificial minichromosome Ch16 expressing ADE6. Progeny cells were selected on YE4S plates lacking adenine. Cells containing the minichromosome were plated onto YE4S plates lacking adenine and incubated at 30ᵒC for 3 days. Cells retaining the minichromosome appeared white, while those losing the minichromosome appeared red.

## Acknowledgments

We thank Vincent Fraisier and Chloe Guedj for the maintenance of microscopes at the PICT-IBiSA Imaging facility (Institut Curie), a member of the France-BioImaging national research infrastructure. We thank Anne Houdusse (Institut Curie), Sergio Rincon (University of Salamanca), Lara Kruger (Cambridge University), and Manuel Lera-Ramirez (University College London) for helpful suggestions on this work. Priyanka Sasmal is supported by a PhD fellowship from the Institut Curie International PhD – EURECA program (Marie Skłodowska-Curie Grant No 847718) and a 4^th^ year PhD fellowship from the Fondation ARC. This work is supported by grants from the Fondation ARC and La Ligue National Contre le Cancer. The Tran team is a member of the Labex CelTisPhyBio, part of IdEx PSL.

## Author Contributions

Priyanka Sasmal: Conception / Experimentations / Analysis / Writing Makito Miyazaki: Conception / Experimentation / Editing Frederique Carlier-Grynkorn: Experimentation Phong Tran: Conception / Writing / Editing / Funding

---

**Figure 1S.**
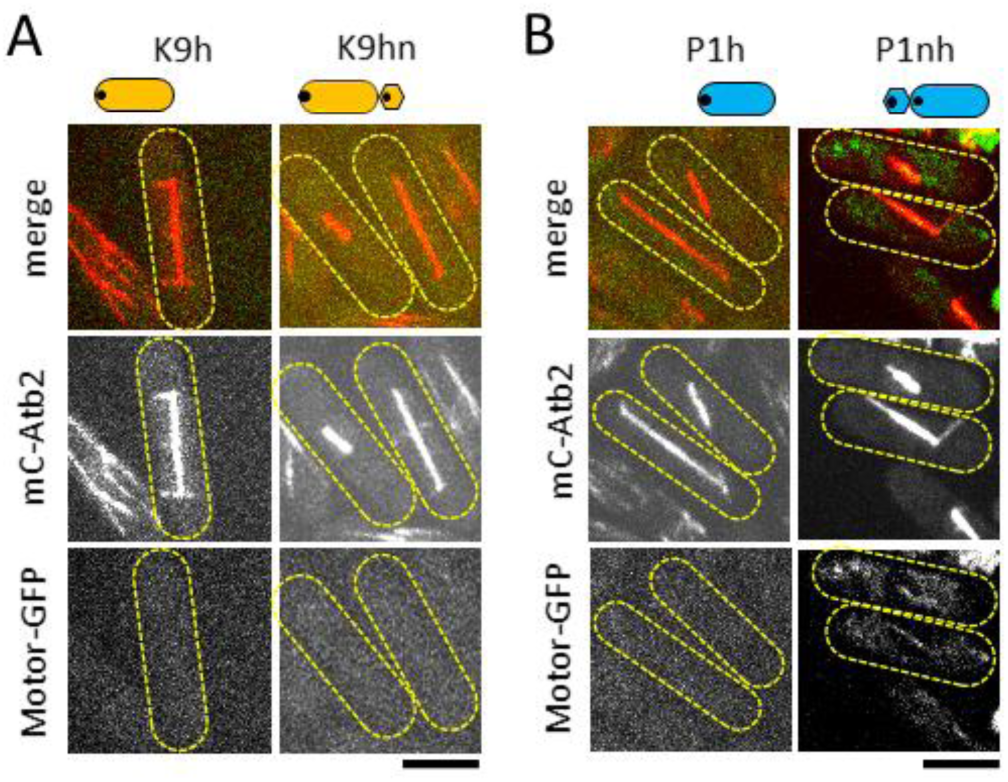
Supplement to Figure 1. **A.** Images of the Klp9 series. Merge color and individual black-white fluorescence of mCherry-Atb2 (tubulin) and motor-eGFP (motors), of mitotic cells (yellow dashed outline) with short metaphase spindles and long anaphase spindles. The chimeric and truncated motors are labelled as cartoons and text. Scale bar, 5 µm. **B.** Images of the Pkl1 series. Merge color and individual black-white fluorescence of mCherry-Atb2 (tubulin) and motor-eGFP (motors), of mitotic cells (yellow dashed outline) with short metaphase spindles and long anaphase spindles. The chimeric and truncated motors are labelled as cartoons and text. Scale bar, 5 µm.

**Figure 2S.**
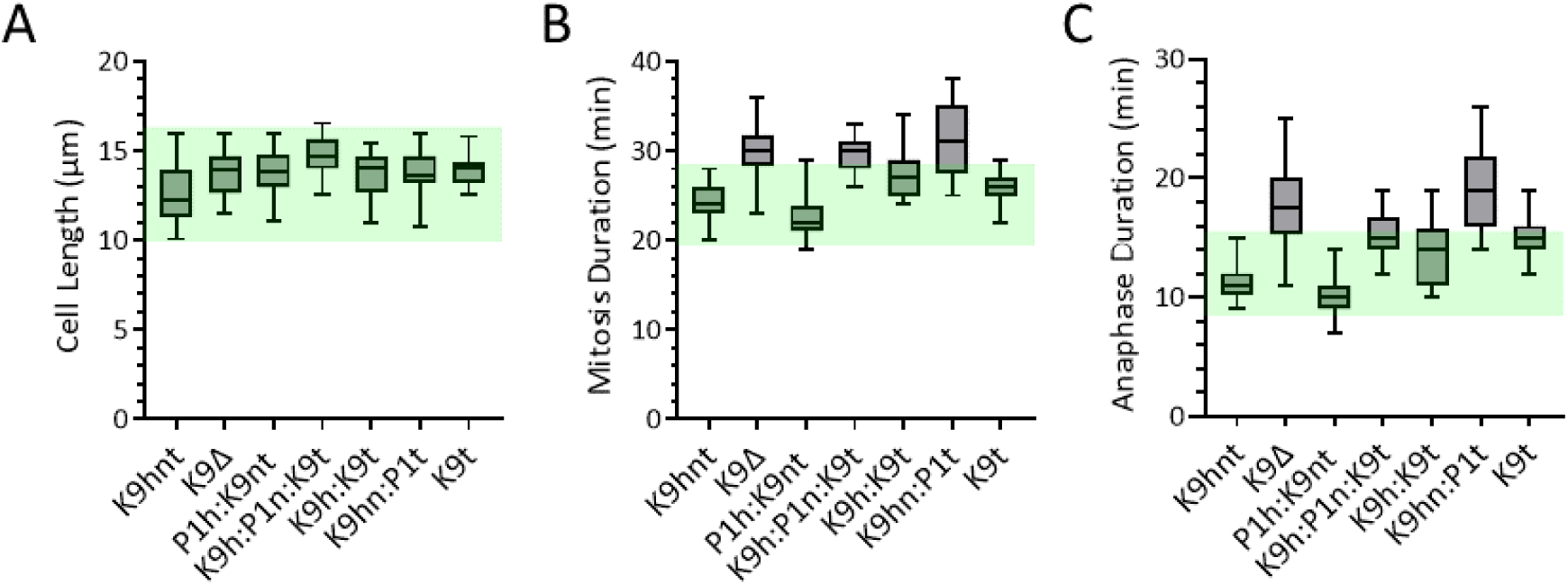
Supplement to Figure 2. **A.** Box plot of Cell Length at mitosis for the Klp9 series. Mean ± Standard Deviation, t-test p-values with respect to wild-type: K9hnt 12.59 ± 1.57 µm (n=20); K9Δ 13.78 ± 1.29 µm (n=20, p<0.05); P1h:K9nt 13.86 ± 1.26 µm (n=20, p<0.05); K9h:P1n:K9t 14.72 ± 1.13 µm (n=20, p<0.001); K9h:K9t 13.62 ± 1.33 µm (n=20, p<0.05); K9hn:P1t 13.91 ± 1.18 µm (n=20, p<0.01); K9t 14.02 ± 0.87 µm/min (n=20, p<0.005). **B.** Box plot of total Mitosis Duration for the Klp9 series. Mean ± Standard Deviation, t-test p-values with respect to wild-type: K9hnt 24 ± 2 min (n=20); K9Δ 30 ± 3 min (n=20, p<0.001); P1h:K9nt 23 ± 3 min (n=20, p<0.001); K9h:P1n:K9t 29 ± 2 min (n=20, p<0.001); K9h:K9t 28 ± 3 min (n=20, p<0.001); K9hn:P1t 32 ± 4 min (n=20, p<0.001); K9t 26 ± 2 min (n=20, p<0.05). **C.** Box plot of Anaphase B Duration for the Klp9 series. Mean ± Standard Deviation, t-test p-values with respect to wild-type: K9hnt 11 ± 2 min (n=20); K9Δ 18 ± 4 min (n=20, p<0.001); P1h:K9nt 10 ± 2 min (n=20, p<0.05); K9h:P1n:K9t 15 ± 2 min (n=20, p<0.001); K9h:K9t 14 ± 3 min (n=20, p<0.005); K9hn:P1t 19 ± 4 min (n=20, p<0.001); K9t 15 ± 2 min (n=20, p<0.001).

**Figure 3S.**
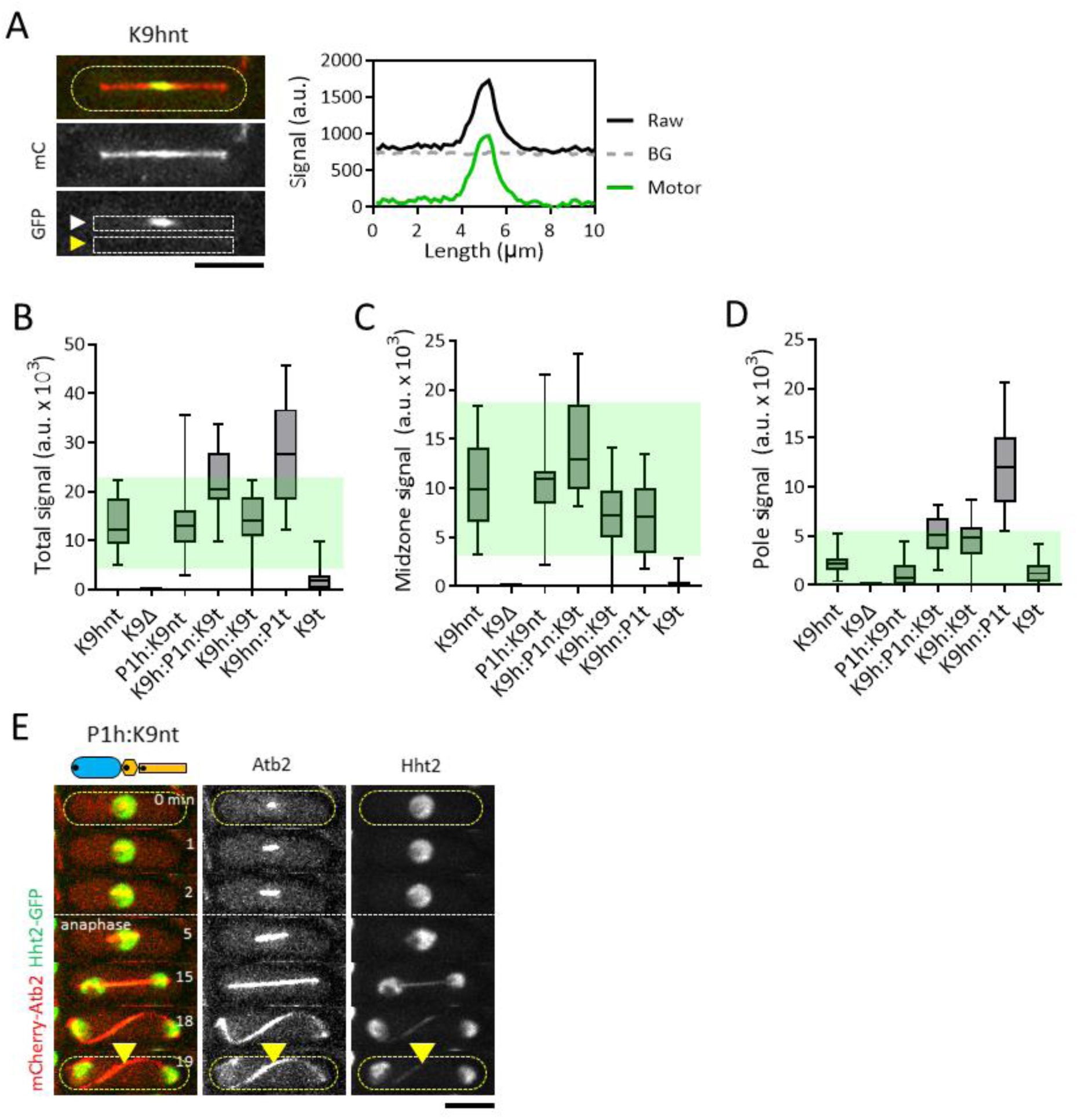
Supplement to Figure 3. **A.** Line scan method to measure motor-GFP signal intensity on the 10-µm long spindle. White arrow head indicates line scan of Raw signal. Yellow arrow head indicates line scan of Back Ground signal. Plot of Signal versus Length shows Raw, Back Ground (BG), and Motor (Raw – BG) signal. **B.** Box plot of Total Signal intensity of motor-GFP on 10-µm long spindles of the Klp9 series. Signal is in arbitrary unit (a.u. x 10^3^). Mean ± Standard Deviation, t-test p-values with respect to wild-type: K9hnt 13.2 ± 5.3 (n=20); K9Δ 0 ± 0 (n=20); P1h:K9nt 13.9 ± 7.0 (n=20, p=0.7); K9h:P1n:K9t 21.8 ± 6.2 (n=20, p<0.001); K9h:K9t 14.4 ± 5.4 (n=20, p=0.5); K9hn:P1t 27.6 ± 10.0 (n=20, p<0.001); K9t 2.3 ± 2.6 (n=20, p<0.001). **C.** Box plot of Midzone Signal intensity of motor-GFP on 10-µm long spindles of the Klp9 series. Signal is in arbitrary unit (a.u. x 10^3^). Mean ± Standard Deviation, t-test p-values with respect to wild-type: K9hnt 10.4 ± 4.5 (n=20); K9Δ 0 ± 0 (n=20); P1h:K9nt 10.6 ± 4.2 (n=20, p=0.9); K9h:P1n:K9t 14.0 ± 4.6 (n=20, p<0.05); K9h:K9t 7.5 ± 3.6 (n=20, p<0.05); K9hn:P1t 7.1 ± 3.8 (n=20, p<0.05); K9t 0.5 ± 0.8 (n=20, p<0.001). **D.** Box plot of combined spindle Pole Signal intensity of motor-GFP on 10-µm long spindles of the Klp9 series. Signal is in arbitrary unit (a.u. x 10^3^). Mean ± Standard Deviation, t-test p-values with respect to wild-type: K9hnt 2.2 ± 1.1 (n=20); K9Δ 0 ± 0 (n=20); P1h:K9nt 1.3 ± 1.4 (n=20, p<0.05); K9h:P1n:K9t 5.1 ± 1.8 (n=20, p<0.001); K9h:K9t 4.5 ± 2.1 (n=20, p<0.001); K9hn:P1t 12.4 ± 4.5 (n=20, p<0.001); K9t 1.4 ± 1.1 (n=20, p<0.05). **E.** Time-lapsed images of spindle dynamics throughout mitosis for chimera P1h:K9nt. Cell expressed mCherry-Atb2 (tubulin) and Hht2-GFP (histone). Yellow arrow head indicates medial position where cell divides. Scale bar, 5 µm.

**Figure 4S.**
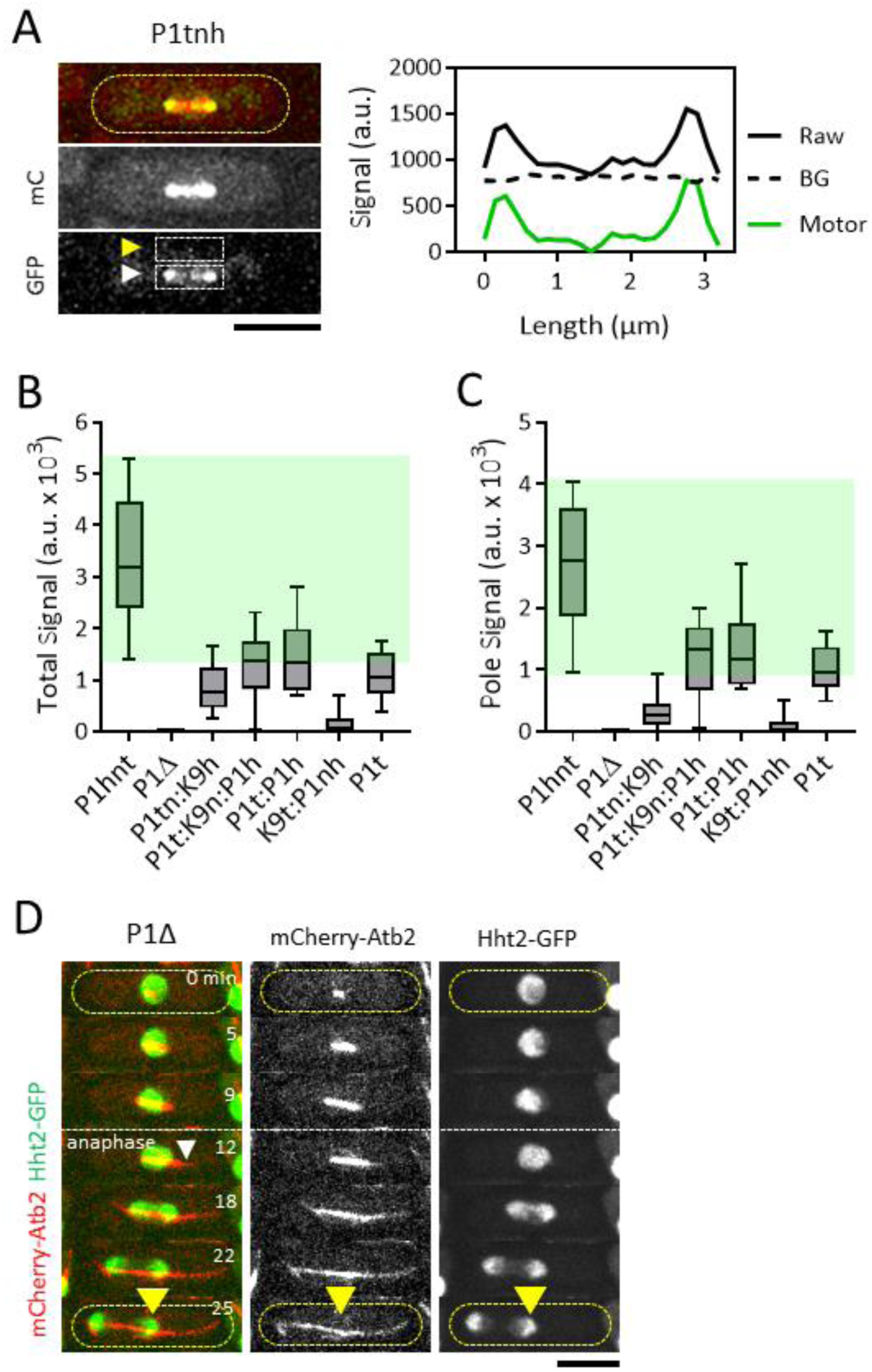
Supplement to Figure 4. **A.** Line scan method to measure motor-GFP signal intensity on the 3-µm long spindle. White arrow head indicates line scan of Raw signal. Yellow arrow head indicates line scan of Back Ground signal. Plot of Signal versus Length shows Raw, Back Ground (BG), and Motor (Raw – BG) signal. **B.** Box plot of Total Signal intensity of motor-GFP on 3-µm long spindles of the Pkl1 series. Signal is in arbitrary unit (a.u. x 10^3^). Mean ± Standard Deviation, t-test p-values with respect to wild-type: P1tnh 3.35 ± 1.29 (n=20); P1Δ 0 ± 0 (n=20); P1tn:K9h 0.84 ± 0.44 (n=20, p<0.001); P1t:K9n:P1h 1.28 ± 0.69 (n=10, p<0.001); P1t:P1h 1.43 ± 6.7 (n=10, p<0.001); K9t:P1nh 0.17 ± 0.27 (n=10, p<0.001); P1t 1.09 ± 0.44 (n=10, p<0.001). **C.** Box plot of combined Pole Signal intensity of motor-GFP on 3-µm long spindles of the Pkl1 series. Signal is in arbitrary unit (a.u. x 10^3^). Mean ± Standard Deviation, t-test p-values with respect to wild-type: P1tnh 2.70 ± 1.04 (n=20); P1Δ 0 ± 0 (n=20); P1tn:K9h 0.31 ± 0.24 (n=20, p<0.001); P1t:K9n:P1h 1.17 ± 0.63 (n=10, p<0.001); P1t:P1h 1.31 ± 6.3 (n=10, p<0.001); K9t:P1nh 0.09 ± 0.16 (n=10, p<0.001); P1t 1.02 ± 0.37 (n=10, p<0.001). **D.** Time-lapsed images of spindle dynamics throughout mitosis for P1Δ. Cell expressed mCherry-Atb2 (tubulin) and Hht2-GFP (histone). White arrow head indicates a MT protrusion (time = 12 min) that pushed the chromosome mass to the cell center, where subsequent cell division (yellow arrow head) resulted in aneuploidy. Scale bar, 5 µm.

## Supplemental Data (Strain List)

TP 472 h+ miniChromosomeCh16-ADE6ade6-210 leu1-32 ura4-D18

TP 4994 h-Klp9hnt-eGFP:KanR mCherry-Atb2:HygR ade6-M216 leu1-32 ura4-D18

TP 4995 h+ Klp9hnt-eGFP:KanR mCherry-Atb2:HygR ade6-M216 leu1-32 ura4-D18

TP 5015 h-Klp9Δ:NatR mCherry-Atb2:HygR ade6-M216 leu1-32 ura4-D18

TP 5016 h+ Klp9Δ:NatR mCherry-Atb2:HygR ade6-M216 leu1-32 ura4-D18

TP 5328 h-Pkl1h:Klp9nt-eGFP:KanR mCherry-Atb2:HygR ade6-M216 leu1-32 ura4-D18

TP 5632 h-Pkl1Δ:Ura4+ mCherry-Atb2:HygR ade6-M210 leu1-32 ura6-D18

TP 5963 h-Klp9t-eGFP:KanR mCherry-Atb2:HygR

TP 5988 h- Pkl1tnh-eGFP:KanR mCherry-Atb2:HygR ade-6-M210 leu1-32 ura6-D18

TP 5989 h+ Klp9h:Klp9t-eGFP:KanR mCherry-Atb2:HygR ade-6 M216 leu1-32 ura-D18

TP 5994 h- Pkl1t:Pkl1h-eGFP:KanR mCherry-Atb2:HygR ade-6-M210 leu1-32 ura6-D18

TP 6013 h+ Klp9h:Pkl1n:Klp9t-eGFP:KanR mCherry-Atb2:HygR ade6-M210 leu1-32 ura4-D18

TP 6016 h+ Klp9hn:Pkl1t-eGFP:KanR mCherry-Atb2:HygR ade6-M210 leu1-32 ura4-D18

TP 6018 h- Klp9t:Pkl1nh-eGFP:KanR mCherry-Atb2:HygR ade6-M210 leu1-32 ura4-D18

TP 6024 h- Pkl1h:Klp9n:Pkl1t-eGFP:KanR mCherry-Atb2:HygR ade-6-M210 leu-32 ura-D18

TP 6030 h- Pkl1t-eGFP:KanR mCherry-Atb2:HygR ade6-M210 leu1-32 ura4-D18

TP 6038 h- Klp9h-eGFP:KanR mCherry-Atb2:HygR ade6-M216 leu1-32 ura4-D18

TP 6039 h+ Klp9hn-eGFP:KanR mCherry-Atb2:HygR ade6-M216 leu1-32 ura4-D18

TP 6040 h- Pkl1h-eGFP:KanR mCherry-Atb2:HygR ade6-M216 leu1-32 ura4-D18

TP 6041 h- Pkl1nh-eGFP:KanR mCherry-Atb2:HygR ade6-M216 leu1-32 ura4-D18

TP 6057 h- Pkl1tn:Klp9h-eGFP:KanR mCherry-Atb2:HygR ade6-M210 leu1-32 ura4-D18

